# Spatial autocorrelation dimension as a potential determinant for the temporal persistence of human atrial and ventricular fibrillation

**DOI:** 10.1101/2023.04.12.536515

**Authors:** Dhani Dharmaprani, Evan V. Jenkins, Jing X. Quah, Kathryn Tiver, Lewis Mitchell, Matthew Tung, Waheed Ahmad, Nik Stoyanov, Martin Aguilar, Martyn P. Nash, Richard H. Clayton, Stanley Nattel, Anand N. Ganesan

## Abstract

**Background:** Despite being central to atrial fibrillation (AF) and ventricular fibrillation (VF) mechanisms and therapy, the factors governing AF and VF termination are poorly understood. It has been noted that ratio of system size (*L*) and the two-point spatial correlation length (ξ_2_) are associated with time until termination in transient spatiotemporally chaotic systems, but the relationship between these characteristics and termination has not been systematically studied in human AF and VF.

**Objective:** We aimed assess whether the time to cardiac fibrillation termination can be predicted using a novel estimator, the spatial autocorrelation dimension (*D*_*i*_), defined as the ratio of *L* and ξ_2_, in human AF and VF.

**Methods:** *D*_*i*_ was computed and compared in a multi-centre, multi-system study with data for sustained versus spontaneously terminating human AF/VF. VF data was collected during coronary-bypass surgery; and AF data during clinically indicated AF ablation. We analyzed: i) VF mapped using a 256-electrode epicardial sock (n=12pts); ii) AF mapped using a 64-electrode constellation basket-catheter (n=15pts); iii) AF mapped using a 16-electrode HD-grid catheter (n=42pts). To investigate temporal fibrillation persistence, the response of AF-episodes to flecainide (n=7pts) was also studied.

**Results:** Spontaneously terminating fibrillation demonstrated a lower *D*_*i*_ (P<0.001 all systems). Lower *D*_*i*_ was also seen in paroxysmal compared to persistent AF (P=0.002). Post-flecainide, *D*_*i*_ decreased over time (P<0.001). Lower *D*_*i*_ was also associated with longer-lasting episodes of AF/VF (R^2^>0.90, P<0.05 in all cases). Using k-means clustering, two distinct clusters and their centroids were identified i) a cluster of spontaneously terminating episodes, and ii) a cluster of sustained epochs.

**Conclusion:** *D*_*i*_ predicts the temporal persistence of cardiac fibrillation. This finding provides potentially important insights into a possible common pathway to termination and therapeutic approaches.

## INTRODUCTION

Cardiac fibrillation is a spatiotemporally disordered electromechanical state observed in either the atria or ventricles.^1^ A fundamental property of the arrhythmia is the propensity for some episodes of fibrillation to spontaneously terminate and transition into sinus rhythm, while other episodes may persist. In the case of atrial fibrillation (AF), individual episodes may be as short as a few seconds, but may also last for hours, days or become permanent in individual patients.^2^ Ventricular fibrillation (VF) episodes may also have the propensity VF to self-terminate. However, if VF persists for longer than 10 minutes without resuscitation, the arrhythmia can lead to irreversible neurologic harm or death.^3, 4^

The general properties of fibrillatory termination and persistence are of particular clinical interest as they have the potential to inform ongoing efforts to achieve personalized prediction of prognosis and therapy in both human fibrillatory disorders. Although extensive efforts have occurred over many decades to empirically identify predictors of termination or persistence in cardiac fibrillation,^5, 6^ at present there is no universally accepted conceptual paradigm or metric to explain why individual episodes of fibrillation may have the propensity to persist or spontaneously terminate. Such a general paradigm, if it could be developed, would have significant practical relevance, as it could potentially inform unresolved contemporary clinical questions. Examples of such challenges could include the persistence of AF despite pulmonary vein isolation^7^, the upper limit of effectiveness observed with PVI,^7^ objective delineation of paroxysmal from persistent forms of AF,^8^ or understanding why some episodes of VF are responsive or unresponsive to defibrillation shocks.^9^

Here, we sought to contribute to efforts to develop a conceptual paradigm, to account for the potential for episodes of fibrillation to either persist or self-terminate. The electromechanical complexity that characterizes AF and VF has frequently been compared to the sustained disorder of nonequilibrium chaotic systems in nature that have been observed in physical,^10, 11^ chemical^12^ and biological systems.^13^. An important general property observed in these systems is synchronization of behavior between spatially connected neighboring regions, that gradually breaks down with distance, a property that can be measured with the two-point spatial correlation length (ξ_2_).

In an early study of spatiotemporal chaotic systems by Egolf and Greenside, it was postulated that the ratio of system size (*L*) to correlation length (ξ_2_), along with knowledge of spatial dimensionality (*d*) could be used to decompose the behavior of chaotic systems into ‘independent regions.’ ^14^ Here, we name this quantity, which was not formally defined in the original paper, as the spatial autocorrelation dimension (*D*_*i*_). The spatial autocorrelation dimension is defined with the following equation:

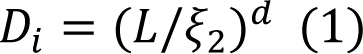

where *L* the relative system size, ξ_2_ the two-point correlation between pairwise electrodes spatially spaced Δ*x* apart (i.e. inter-electrode spacing), and *d* the spatial dimensionality of the system. As an example of how the *D*_*i*_ estimator could potentially be computed, Egolf and Greenside suggested the following estimator for a heart with a radius *R* and *d* = 2 (2):

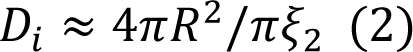

To our knowledge, the association of the spatial autocorrelation dimension estimator with the persistence of termination with human cardiac fibrillation has never formally been assessed. In this study, we evaluate the hypothesis that differences in *D*_*i*_ may account for the relative time until termination in AF and VF.^14, 15^ We explore this by calculating *D*_*i*_ in spontaneously-terminating versus sustained episodes of AF and VF, as well as in episodes of AF post pharmacological intervention, with supporting data from computer simulations.

## MATERIALS & METHODS

### Epicardial sock recordings of human VF

The human VF study uses prospectively collected data as described by Nash et al.^16^ The study recruited n=12 patients (n=8 sustained VF, n=4 self-terminating VF) undergoing routine coronary bypass graft procedures for ischemic heart disease with cross-clamp fibrillation. These studies were approved by the Hospital Ethics Committee (REC 01/0130) with written informed consent. Individual patient details are given in **Supplementary Materials S1, Supplemental Table 1**. Recordings were obtained using a 256-electrode epicardial sock (Δ*x* = 10mm inter-electrode spacing) and unipolar electrograms (1kHz sampling) were preprocessed as previously described.^17, 18^ In a subset of patients, VF spontaneously terminated before the full 210 seconds without requiring defibrillation, defined as ‘self-terminating episodes’. This was compared to sustained VF episodes lasting the full duration. Additional study details are provided in **Supplement S1**.

### Basket and HD grid recordings of human AF

The human AF study extends our previously published study.^19^ The study was a multi-center observational design analyzing prospectively collected electrophysiologic data acquired prior to ablation (HREC110634). The inclusion criterion was AF undergoing ablation. Patient participation was by informed consent, with recruitment from Flinders University and Hamburg University. The patient cohort included n=15 patients (age 62±8, 13/15 male(87%), BMI 28.7±3.8 kg/m^2^, non-paroxysmal 12/15(80%); **S1, Supplemental Table 2**). Basket catheter and HD-grid recordings were performed as previously.^20^ 64-electrode basket catheters (Constellation, Boston Scientific, 48mm (Δ*x* = 4mm), 60mm (Δ*x* = 5mm) and HD-grid a 3-3-3 mm catheter with 16 electrodes to record human AF (Δ*x* = 3mm) were utilized to record unipolar electrograms (1-500 Hz, 2000Hz sampling) and surface electrocardiograms during spontaneous or induced AF lasting >1 minutes. Spontaneously terminating AF episodes were defined as those that terminated before the full 1-minute without requiring defibrillation, whilst sustained recordings were defined as those lasting the full window of observation. Signal filtering, cleaning, and processing are detailed further in **Supplement S3**.

### Spatial autocorrelation dimension estimation

To calculate the spatial autocorrelation dimension, *D*_*i*_, atrial and ventricular volumes were first calculated using the convex hull of electrode co-ordinates from one of the following i) mapping catheter (for atrial fibrillation), ii) epicardial sock (ventricular fibrillation) or iii) grid nodes (computer simulations). This volume was then used to estimate the radius, *R*, which was related to the two-point correlation, ξ_2_, to give the spatial cross-correlation as given by equation (2) (a graphical overview of *D*_*i*_ calculation and further details are provided in **Extended Data Figure 1, Supplement S3**).^21^ Specifically, ξ_2_ gives the association between the maximum cross correlation and distance between two pairwise signals recorded at different electrodes, which are spatially distanced Δ*x* apart. This can often be modelled by an exponential relationship^22^ (details in **Supplement S3**):

**Figure 1:**
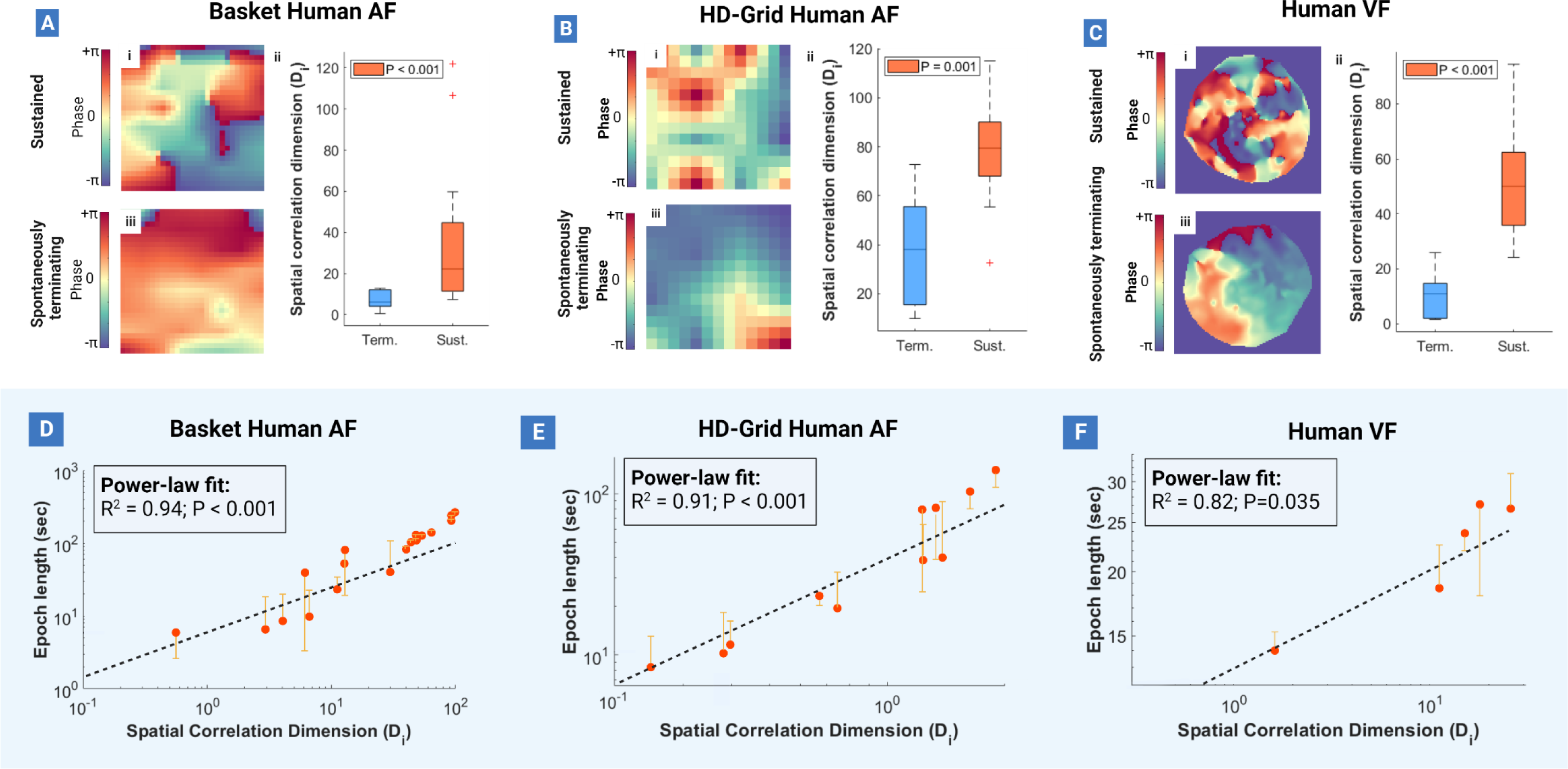
Spatial autocorrelation dimension reflects the likelihood of AF/VF termination and epoch length. (**1A-C**): Terminating epochs of AF and VF are associated with lower spatial autocorrelation dimensions (1A-Cii). Observed epoc lengths (which gives time until termination) increased with larger spatial autocorrelation dimensions, with a relationship that best fit to a power-law (**1D-F**).

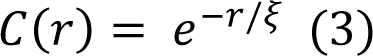

The cross-correlation function between two sequences of normalized activations were calculated over windows shifted throughout the duration of the recording, with a range of lag values encompassing slightly more than one cycle length (∼170ms for VF and ∼150ms for AF). The maximum cross-correlation was given by the correlation coefficient for the sequence of activations between the two corresponding locations. This process was repeated in a pairwise fashion to calculate the correlation between all possible combinations of locations, with locations corresponding to electrodes for AF and VF data, or mesh vertices for computer simulations.

### Assessing the effect of pharmacological intervention

To understand the effect by which anti-arrhythmic medicines facilitate termination of fibrillation, we studied the action of prototype sodium-channel blockade as archetypal pharmacologic rhythm modulation strategy on spatial autocorrelation dimension. This was assessed in: i) n=9 computer simulated epochs of AF with varying levels of I_Na_ inhibition (20%-60%) leading to AF termination; and ii) n=21 epochs from 7 AF patients administered flecainide during an electrophysiology study, mapped using the HD-grid catheter.

To simulate the effect of sodium-channel block in computer simulated AF, the two-dimensional Courtemanche-Ramirez-Nattel cell model was implemented as previously described (run on 7×6cm square grids).^23^ Differential equations were solved with a 25ms time-step, with simulations run for 5 seconds. AF epochs were initiated by a standard S1-S2 cross-shock protocol. To achieve spontaneous AF termination, ACh gradients were altered to a range of values to achieve self-termination before the full 5 seconds (**Supplement S2**).

### Comparison with other metrics characterising arrythmia dynamics

#### Electrogram-based metrics

To evaluate whether *D*_*i*_ provides new information, we evaluated whether established electrogram-based metrics used to characterise arrythmia dynamics would associate with spontaneously terminating AF and VF. This was assessed by calculating four metrics using the HD-grid human AF data: i) Dominant Frequency (DF)^24^, Shannon Entropy (ShEn)^25^, mean bipolar voltage^26^, and peak-peak bipolar voltage. Details on the calculation of these metrics are provided in **Supplement S4**.

#### Renewal rate constants

A common motif of extensive spatiotemporally disordered systems is the continuous creation and annihilation of topological defects known as phase singularities (PS) PS.^20, 27^ The average rates of PS formation (given by λ_*f*_) and PS destruction (λ_*d*_) can be measured using renewal theory based approaches.^18, 20^ We have previously shown that spontaneous termination of AF and VF may be reflected by lower mixing rates compared to sustained AF and VF, governed by the interaction λ_*f*_ and λ_*d*_ as reflected by the second largest eigenvalue modulus (SLEM) of the Markov transition matrix (Supplement S4).

Here, we build on this observation and hypothesized that changes in λ_*f*_, λ_*d*_, the mixing rate should associate with changes in *D*_*i*_. The average rate of phase singularity (PS) formation (given by λ_*f*_) and destruction (λ_*d*_) can be measured by calculating the exponential decay constant from the inter-event time distribution as previously described^19^ (further detailed in **Supplement S6**). The interaction of λ_*f*_ and λ_*d*_ is given by the ‘*mixing rate*’, which represents the time taken to reach the stationary state distribution (**Supplement S6**). In this study, λ_*f*_ and λ_*d*_ were calculated as previously described from pre-processed and Hilbert transformed unipolar electrograms, with the PS detection algorithm further detailed **Supplement S7**. ^18, 20^

### Statistical Analysis

The association of *D*_*i*_ with temporal persistence of fibrillatory episodes was determined using non-linear least squares regression with *D*_*i*_ as the independent variable and time until observed termination as the dependent variable. Differences in spatial autocorrelation dimension, correlation length, λ_*f*_, λ_*d*_ and mixing rates between cases of sustained and spontaneously terminating AF, VF, and computer simulated fibrillation were assessed using generalized linear mixed models (significance at P<0.05) (**Supplement S8**). A similar approach was used to study spatial autocorrelation dimension and correlation length across different stages of VF and post flecainide administration. To study the relationship between λ_*f*_ and λ_*d*_ with spatial autocorrelation dimension and correlation length, a k-means clustering approach was implemented using 10 replicates and a city-block algorithm as the distance metric. Additional details are provided in **Supplement S8**.

## RESULTS

### Spatial autocorrelation dimension reflects the likelihood AF/VF self-termination and observed epoch length

Sustained episodes of basket-catheter mapped human AF were associated with lower spatial autocorrelation dimension (*D*_*i*_) (mean terminating = 7.10,95%CI:4.77,9.44; mean sustained = 34.38,95%CI:16.87,51.88; P<0.001). This was also seen in HD-grid mapped human AF (mean sustained = 78.78,95%CI:63.35,94.21; mean terminating = 9.41,95%CI:4.49,14,33; P=0.001) and human VF (mean terminating = 10.30,95%CI:-1.90,22.50; mean sustained = 52.80,95%CI:27.13,78.48; P<0.001) (**Figure 1A-C**). Higher *D*_*i*_ also associated with increasing epoch lengths that was best fit to a power-law relationship in basket-mapped human AF (R^2^ = 0.940; P<0.001), HD-grid mapped human AF (R^2^ = 0.910; P<0.001), and human VF (R^2^ = 0.817; P=0.035) when compared to a first-degree polynomial (Basket: R^2^ = 0.921; HD: 0.844; VF: R = 0.817; P<0.001 all systems) and exponential relationship (Basket: R^2^ = 0.913; HD: R^2^ = 0.906; VF: R = 0.753; P<0.001 all systems). (**Fig.1**D-F).

### Relationship to the evolution of human atrial and ventricular fibrillation

*D*_*i*_ were also further assessed in 380 epochs arising from n=24 paroxysmal AF patients, and 282 epochs arising from n=18 persistent human AF patients. Epochs from paroxysmal AF patients were associated with consistently lower *D*_*i*_ in paroxysmal AF (31.09,95%CI:25.47,36.70) when compared to persistent AF (57.73,95%CI:43.93,71.52) (P=0.002; **Figure 2A**).

**Figure 2:**
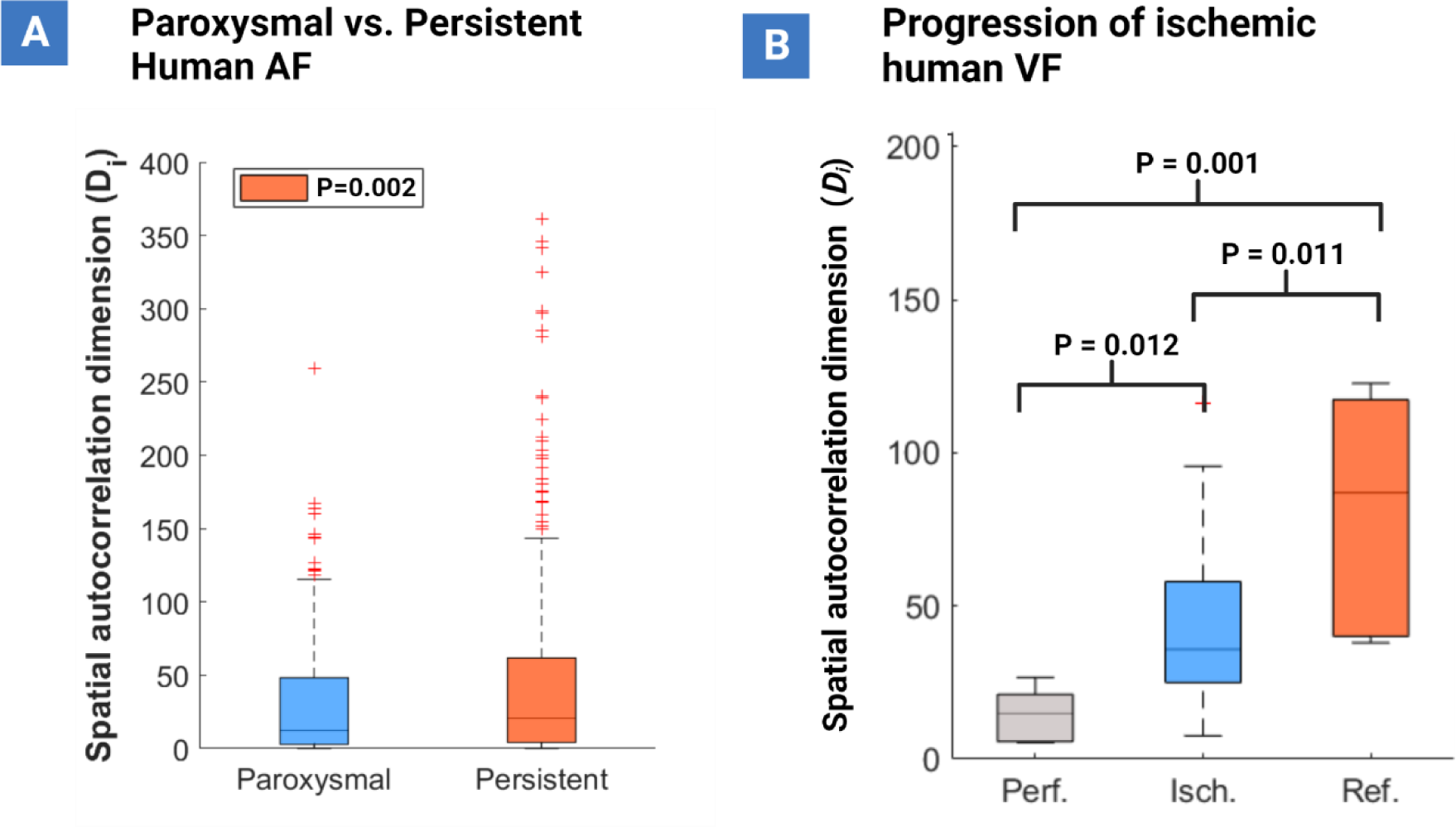
Relationship to the evolution of AF and VF. (**2A**): Paroxysmal AF was associated with lower spatial autocorrelation dimensions compared to persistent AF. (**2B**): As ischemic VF evolves and becomes more turbulent and disordered, spatial autocorrelation dimension increased.

Human ventricular fibrillation episodes from n=12 patients undergoing cardiac surgery were also studied. Three distinct stages of VF were studied: i) the perfusion stage (initial 30 seconds); ii) ischemia (following 150 seconds); and iii) reflow (last 30 seconds). **Extended Data Fig.2, Supplement S9** depicts the propagation of electrical activity as VF evolves throughout the three stages, which can be seen to depict more turbulent behaviour as VF progresses. This evolution was associated with larger *D*_*i*_ (mean perfusion = 14.78,95%CI:5.33,24.67); mean ischemia = 45.48,95%CI:33.36,57.12); mean reflow = 82.06,95%CI:41.78,122.22). Specifically, *D*_*i*_ between the perfusion and ischemia (P=0.012), and perfusion and reflow (P<0.001), and ischemia and reflow stages were significantly different (P=0.011) (**Figure 2B**).

### Relationship to other fibrillatory metrics

#### Rates of PS formation and destruction

In all model systems, the rate of PS formation, λ_*f*_, was lower in spontaneously terminating basket-mapped human AF (terminating = 0.017,95%CI:0.015,0.019); sustained = 0.027,95%CI:0.025,0.030), HD-grid human AF (terminating = 0.0022,95%CI:0.0017,0.0028; sustained = 0.007,95%CI:0.0066,0.0075), and human VF (terminating = 0.011,95%CI:0.010,0.012; sustained = 0.023,95%CI:0.024,0.025). Similar trends were seen for the rate of PS destruction, λ_*d*_ in basket-mapped human AF (terminating = 0.016,95%CI:0.014,0.017; sustained = 0.022,95%CI:0.021,0.23), HD-grid human AF (terminating = 0.026,95%CI:0.021,0.031); sustained = 0.060,95%CI:0.055,0.065), and human VF (terminating = 0.0103,95%CI:0.009,0.0106; sustained = 0.46,95%CI:0.42,0.50).

When related to the spatial autocorrelation dimension and assessed using k-means clustering, two distinct clusters and their centroids were identified: i) a cluster of self-terminating epochs situated towards the upper right-hand side of graph (orange), and ii) a cluster of sustained epochs on the lower left-hand side of the graph (blue) (Fig.3A-C i **and ii**). In all model systems, the two clusters gave large positive silhouette values (**Fig.3A-C iii and iv**), indicating that the clusters are well separated. The changes seen in λ_*f*_ and λ_*d*_ also led to larger SLEMs and slower mixing rates characterizing episodes of spontaneously terminating fibrillation in all model systems (**Fig.4A-D**).

**Figure 3:**
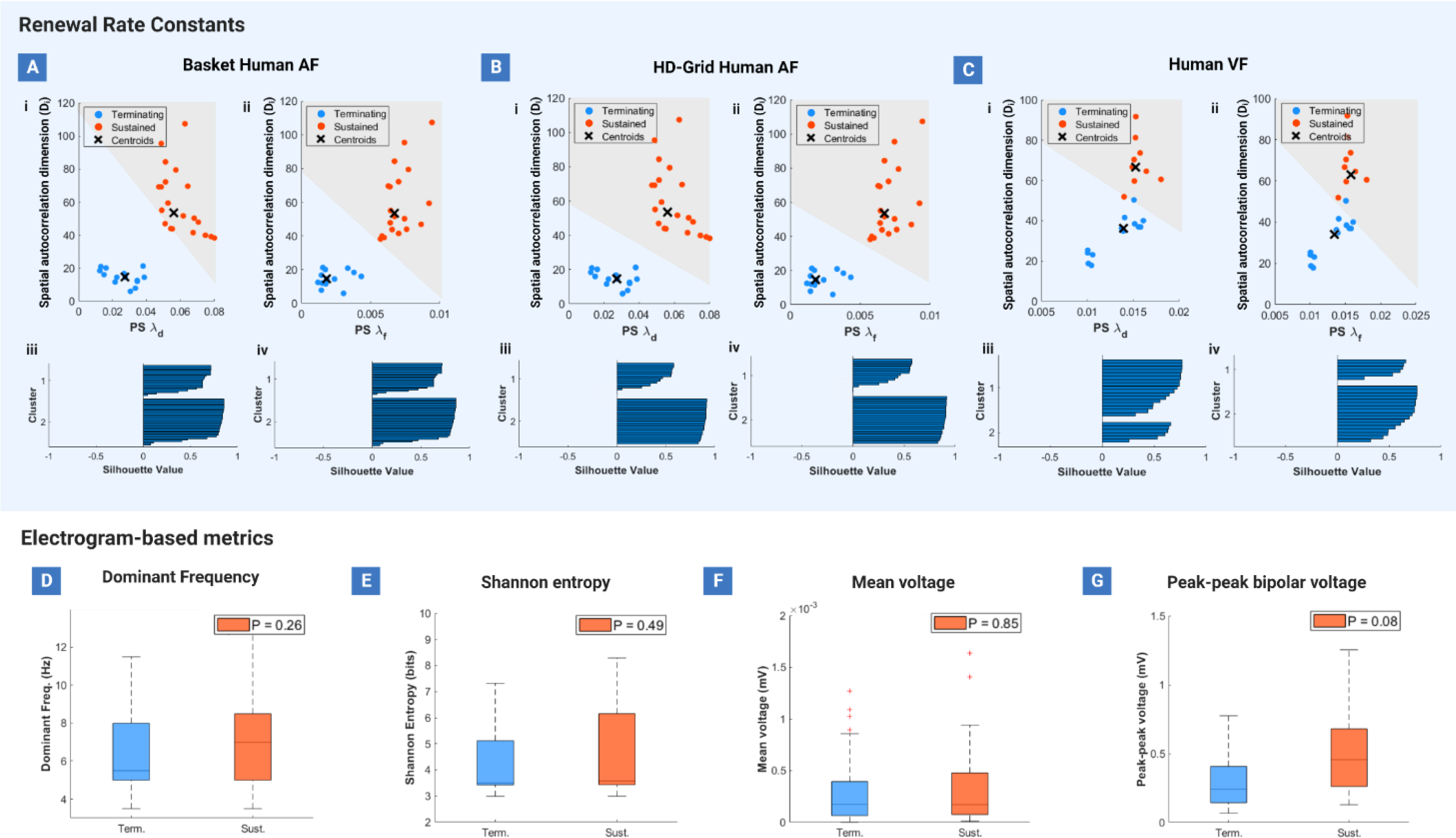
Spatial autocorrelation dimension vs. renewal rates and comparison to other metrics. (**3A-C**): Rates of PS formation (λ_*f*_) and destruction (λ_*d*_) reduced with lower spatial correlation (*D*_*i*_) in terminating epochs of AF and VF. When compared to sustained AF and VF epochs, two clusters were identified by k-means clustering. This separated and spontaneously terminating (blue) and sustained (orange) simulations ((i) and (ii)). The large positive silhouette values indicate good separate of these clusters ((iii) and (iv)). (**3D-G**): Electrogram-based metrics such as dominant frequency (DF), Shannon entropy, mean bipolar voltage, and peak-peak bipolar voltage showed a trend towards lower values in terminating AF/VF, but no statistically significant differences when compared in sustained versus spontaneously terminating AF and VF epochs.

**Figure 4:**
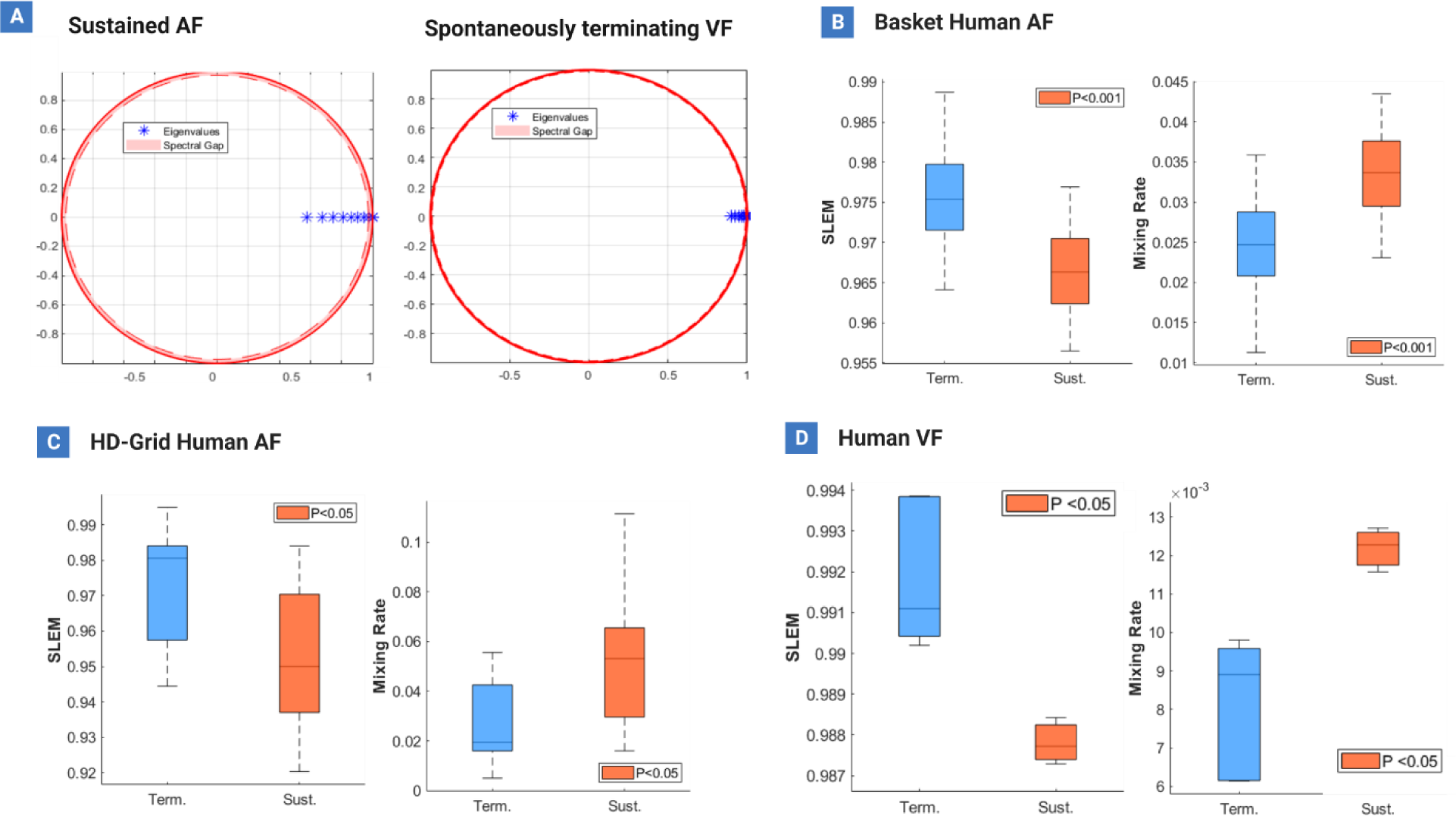
Changes in SLEM and mixing rate. (**4A**): Decreases in λ_*f*_ and λ_*d*_ are reflected by increases in the second largest eigenvalue modulus (SLEM) of the Markov transition matrix. In all model systems, cases of sustained fibrillation demonstrated lower SLEM compared to self-terminating episodes, indicated by the greater spread of eigenvalues on the eigenvalue spectrum plot. (**4B-D**): The higher SLEM seen in self-terminating episodes also lead to a lower mixing rate when compared to sustained episodes in all model systems.

### Electrogram-based measures

When applied to human AF data, it was found that DF showed a trend towards lower values in in spontaneously terminating epochs, however, no statistically significant differences were found when compared to sustained AF epochs (mean sustained = 7.11Hz,95%CI:6.46,7.76; mean terminating = 6.522Hz,95%CI:5.74,7.31; P = 0.26) (**Fig.3D**). Similar observations were made for other electrogram-based metrics, with a downward trend in values associated with terminating episodes, but no statistically significant differences observed for ShEn (mean sustained = 4.59 bits,95%CI:4.02,5.16; mean terminating = 4.31 bits,95%CI:3.85,4.77; P=0.49), mean bipolar voltage (mean sustained = 4.5×10^-4^ mV,95%CI:0.26×10^-3^,0.63×10^-3^; mean terminating = 4.20×10^-4^ mV,95%CI:0.14×10^-3^,0.70×10^-3^; P=0.85), and peak-peak bipolar voltage (mean sustained = 0.57 mV,95%CI:0.41,0.73; mean terminating = 0.28 mV,95%CI:0.22,0.35; P=0.08) (**Fig.3D**).

### Changes in response to anti-arrhythmic drug modulation

In cases of computer simulated AF, generalized linear mixed models identified a decreasing trend in spatial autocorrelation dimension (*D*_*i*_) **(Figure 5A ii**), rate of PS formation (λ_*f*_), **Figure 5A iii**) and PS number (**Figure 5C iv**) with increasing Na+ block, but no significant change was seen for the rate of PS destruction (λ_*d*_). Statistically significant differences were also observed in control versus Na^+^-channel block simulations for *D*_*i*_ (P<0.001) and PS number (P<0.001), but not the PS formation rate λ_*f*_ (P=0.19).

**Figure 5:**
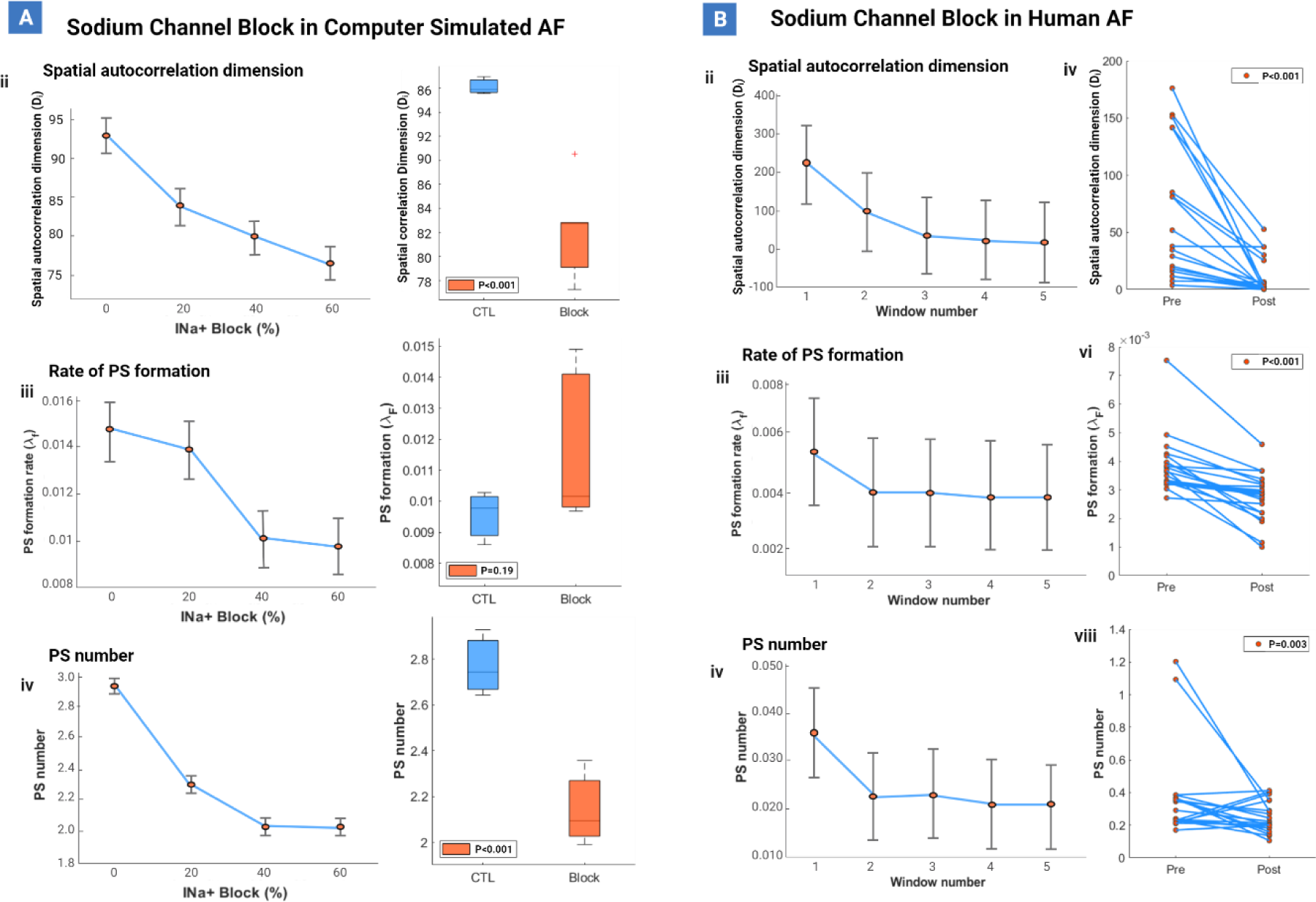
Changes in response to anti-arrhythmic drug modulation. (**5A**): A decrease in spatial autocorrelation dimension *D*_*i*_, λ_*f*_ and PS number was observed over time in Na^+^ block simulations, but not λ_*d*_. (**5B**): In episodes of flecainide-infused human AF, a decrease in spatial autocorrelation dimension *D*_*i*_, λ_*f*_ and PS number was also seen, but not λ_*d*_.

In cases of human AF, the generalized linear mixed model identified a decreasing trend in *D*_*i*_ (**Figure 5B ii**), rate of PS formation (λ_*f*_, **Figure 5B iii**) and PS number (**Figure 5B iv**) in post-flecainide recordings over time, but no significant change in the rate of PS destruction (λ_*d*_). Statistically significant differences were also observed pre-versus post-flecainide for *D*_*i*_ (P<0.001), λ_*f*_ (P<0.001) and PS number (P=0.003).

## DISCUSSION

### Key insights from this study

In this study we aimed to evaluate the hypothesis that spatial autocorrelation dimension (*D*_*i*_) could represent a potentially useful approach to evaluating the potential for persistence or termination in atrial and ventricular fibrillation. In multiple human systems studied, we observed that *D*_*i*_ was consistently higher in episodes that were more sustained (lasting the full period of observation) relative to episodes that self-terminated. This trend was also observed in patients classified using the clinical persistent (lasting >7 days) definition of AF, and further supported in episodes of AF where no anti-arrhythmic drugs were administered. These findings demonstrate that differences in *D*_*i*_ may be a useful tool to characterise the potential for persistence or termination of individual AF and VF episodes. Evidence supporting this hypothesis may offer potentially important clinical and mechanistic implications including: i) provide a quantitative framework to understand the termination of cardiac fibrillation; ii) link termination in both atrial and ventricular to a common mechanism; iii) link the termination of cardiac fibrillation to other nonlinear systems; and iv) inform future investigations of unresolved clinical challenges in human fibrillatory disorders.

### Electrophysiologic characterisation of human AF & VF is an important clinical need

Despite extensive research, the optimal approach to electrophysiologically characterise human fibrillatory dynamics remains incompletely defined. At present, in AF in particular, fundamental challenges remain such as why AF progresses from paroxysmal to persistent in some but not all patients,^8^ why a proportion of patients respond to anti-AF medicines but others do not,^28^ why AF persists despite durable pulmonary vein isolation in some patients,^7, 29^ and why substrate and linear ablation has not led to dramatically improved clinical outcomes.^30^ These problems have led many in the clinical field to suggest that understanding of the biophysical processes sustaining fibrillation is not complete,^31^ providing an important rationale to support the exploration of new strategies for electrophysiologic characterization of human fibrillatory dynamics.

### Spatial autocorrelation dimension may add to available tools to characterise AF and VF dynamics

*D*_*i*_ is an estimator of local electrophysical spatial synchronization in AF and VF, in comparison to overall system size, with the key finding that this metric appears to be a potentially important predictor of the potential for the temporal persistence of fibrillatory episodes. Here, we use the term estimator in the statistical sense, of acting as a rule or criterion for calculating an estimate of a parameter from observable data.^32^

*D*_*i*_ offer a number of potential advantages for the purpose of characterisation of human fibrillatory dynamics, that may make this metric provide useful information in addition to established quantitative tools to explore fibrillatory dynamics. Firstly, the information captured by *D*_*i*_ leverages the inherent spatial properties of the AF and VF. While traditional metrics of fibrillatory dynamics, such as dominant frequency^24^, entropy^25^, voltage^26^ and fractionation^33^ are primarily reliant on local timing and electrogram properties, *D*_*i*_ is an integrated measure over spatially distributed data points, taking into account the overall system size. This may potentially provide important nuanced insights into the system’s underlying dynamics. The spatially informed perspective provided by *D*_*i*_ potentially allow for more accurate consideration of the multi-scale and nonlinear behavior, providing insight into the structural and functional organisation of AF and VF as biophysical processes.

Secondly, *D*_*i*_ is attractive because it is conceptually plausible as potential metric to explain the persistence of fibrillation. In its provenance, *D*_*i*_ has clear theoretical foundations in nonlinear systems theory, although fascinatingly, in the Egolf and Greenside study that motivated this paper *D*_*i*_ was not formally named or defined^14^, despite the hypothesis that this estimator could potentially be used to characterise spatial synchronization in fibrillation being presented as a potential use case.^14^ Mechanistically, *D*_*i*_ is dependent on three factors: (i) the system size; (ii) correlation length; (iii) and spatial dimensionality, analogous to the aspect ratio in extensive chaotic systems.^34^ Furthermore, the foundations for *D*_*i*_ are supported by long-standing empirically derived principles in cardiac fibrillation. The critical mass hypothesis has long been a principle^35 36, 37^ underlying the persistence of fibrillation, and the clinical association of increasing chamber size with the probability of AF maintenance is a generally accepted clinical principle.^36, 37^ Separately, the association of spatial correlation as tool to differentiate paroxysmal from persistent forms of AF has historically been independently clinically validated.^38, 39^ *D*_*i*_, as an integrated estimator of spatial synchronisation relative to system size, has the potential to usefully add to available tools for the electrophysiologic characterization of fibrillation.

Thirdly, a key strength of strength of *D*_*i*_ is that it is potentially able to be measured using clinically available data and has the potentially to leverage recent advances in catheter mapping technologies. Quantification of other similar metrics such as wavelength, has been a challenge for in vivo clinical implementation due to the need for the precise calculation of conduction velocity (CV), effective refractory period (ERP) or action potential duration (APD), difficult to obtain in human patients.^40^ In contrast, *D_i_* can be rapidly calculated from commonly recorded electrical signals such as the intracardiac unipolar & bipolar electrograms, which are routinely collected in electrophysiology studies.

Recent advances in electroanatomic mapping have expanded the potential applications of spatial synchronisation metrics. Novel array catheters^41–43^ allow simultaneous measurement of electrical synchronisation spatial dispersed electrogram information in a variety of potential configurations, which was historically limited in the era of linear mapping catheters.^38^ Additionally, advances in 3D electroanatomic mapping allow rapid and accurate volumetric estimation of cardiac chamber dimensions, allowing system size to be determined.^44^ With new systems integrating real-time export of electrogram information, measurement of *D*_*i*_ has the potential to be clinically available on existing commercial mapping platforms.

It is important to note that because *D*_*i*_ is dependent on system size, its numerical value will vary due to differences in measurement. However, it is interesting to note that *D*_*i*_ estimates were within the same order of magnitude across all model systems. These estimates were also comparable to the number of independent regions originally predicted by Egolf paper et. al. for the fibrillating rabbit heart (∼70).^14^ Another important point to note is that the relationship between *D*_*i*_ and the observed time until AF and VF termination (epoch length) best fit to a power-law, potentially indicating that *D*_*i*_ more accurately predicts shorter lasting episodes of fibrillation with a higher likelihood of impending termination, as opposed to more temporally persistent episodes where termination will occur much later. Therefore, *D*_*i*_ may be clinically useful in applications such as stratifying episodes of AF/VF likely to resolve on their own in the short-term, versus those likely to persist for some time.

Clinically, the spatial autocorrelation dimension could provide a valuable additional quantitative metric to those currently available to measure fibrillatory dynamics.^39, 40^ Firstly, a potential benefit of this approach is that it could contribute to an overarching explanation of why cardiac fibrillation self-terminates. Existing contemporary clinical theories of fibrillation, such as the multiple wavelet re-entry, rotor, or focal source theories of AF and VF^45–47^do not specifically quantitatively address why any individual episode of should have a propensity for self-termination. Similarly, existing quantitative metrics of fibrillatory dynamics such as dominant frequency, entropy, fractionation, or voltage-based substrate characterisation, also leave a potential gap as to why individual epochs of AF and VF self-terminate.^48^

### Links to spatiotemporal turbulence in other systems

An important point to mention is that *D*_*i*_ as presented here to quantify the propensity of AF and VF terminate is linked to concepts that have been extensively explored in theoretical models that have been used to study transient spatiotemporal chaos.^49, 50^ The termination of such transient spatiotemporal chaos has been proposed to occur due to underlying differences in the system phase space, which describes how a system tends to evolve under a variety of initial conditions.^51^ For example, transient spatiotemporal turbulence occurs when a *‘chaotic saddle’* or ‘*non-attracting chaotic set’* is present, which allows turbulent behaviour to eventually terminate when the system escapes into the asymptotic attractor.^15, 52^ *D_i_* represents an index of complexity that approximates the degree of turbulence and number of independent regions in the system^53^, which potentially relates to the likelihood of the system escaping into such an asymptotic attractor.

### Translational perspective

The results presented here may be conceptually useful for understanding and predicting the propensity for the persistence of fibrillation in a variety of clinical contexts. Clinical studies are underway to validate the role of cardiac fibrillation in objectively determining the association of clinically measured *D*_*i*_ with atrial fibrillation persistence, and clinical response to ablation.^54^ *D*_*i*_ has the potential to contribute to the development of new metrics to predict anti-arrhythmic drug response. Clinical studies have recently been commenced to determine the association of *D*_*i*_ and the long-term efficacy of anti-AF treatment. *D*_*i*_ may also have a potentially important role in determining response to AF ablation. An important incompletely resolved clinical question is the role of what if further substrate or linear compartmentalization should occur in patients beyond pulmonary vein isolation.^18^ It is possible that cardiac *D*_*i*_ could play a role in determining which patients could require further ablation beyond PVI, and used as an objective metric to guide clinical responses. Future investigations are underway to answer these questions.

## CONCLUSION

Lower spatial autocorrelation dimension, *D*_*i*_, characterizes the termination of fibrillation. This observation could provide insights into the factors governing termination and persistence of fibrillation and could allow for the development of improved therapeutic approaches based on spatial autocorrelation dimension modulation

## Acknowledgements and funding

We appreciate the comments from Prof. Emeritus Henry Greenside of Duke University, North Carolina, that helped to improve our paper. This work is supported by the National Health and Medical Research Council of Australia Project Grant (1063754) and National Heart Foundation of Australia (101188).

