## Supplementary Materials for "Spatial autocorrelation dimension as a potential determinant for the temporal persistence of human atrial and ventricular fibrillation"

#### S1 – Patient Baseline Demographics

| Patient | Age/Sex | CAD/AVD/MVD | LAD | Cx | RCA | EF, % | MI | B-blockade | ACE | Others |
|---|---|---|---|---|---|---|---|---|---|---|
| **H028** | 77/M | AVD | N | N | N | 74 | No | No | No | No |
| **H034** | 79/M | CAD | Mod | S | N | - | No | Atenolol | Ramipril | No |
| **H048** | 78/M | CAD | N | N | N | 60 | No | No | No | No |
| **H050** | 60/F | CAD | N | N | N | - | No | No | No | No |

Supplemental Table 1: VF patient characteristics

#### S1.2 – Additional VF study details

During the procedure, cardiopulmonary bypass was instituted, and VF induced using 50Hz burst pacing. 30 seconds of control VF was recorded with myocardial perfusion, and the aorta then cross-clamped to achieve global myocardial ischemia. After 150 seconds, the cross-clamp was removed to allow coronary reflow, and a further 30 seconds recorded before defibrillation. Recordings were obtained using a 256-electrode epicardial sock (interelectrode spacing-10 mm).[19] Unipolar electrograms (1kHz sampling) were preprocessed as previously described.[19,20] In a subset of patients, VF spontaneously terminated before the full 210 seconds without requiring defibrillation. These recordings were defined as '*spontaneously terminating episodes*' and compared to sustained VF episodes lasting the full duration.

| Baseline Demographics | All patients (n=15) |
|---|---|
| **Age (years)** | 62±9 |
| **Male, n (%)** | 13 (87) |
| **BMI (kg/m$^2$)** | 28.7±3.8 |
| **$CHA_2DS_2VaSC$** | 1.6±1 |
| **Persistent AF, n (%)** | 12 (80) |
| **Paroxysmal AF, n (%)** | 3 (20) |
| ***Echocardiographic parameters*** | |
| **LVEF (%)** | 59±7 |
| **LA area (cm$^2$)** | 25±4 |
| **E/E’** | 8.4±1.6 |

Supplemental Table 2: Basket-mapped AF Patient Characteristics

| *Baseline Demographics* | *All patients (n=42)* |
|---|---|
| **Age (years)** | 59.1 (9.4) |
| **BMI (kg/m$^2$)** | 31.2 (4.4) |
| **Sex, female [%]** | 12 [28.5] |
| **CHA2DS2-VASc score** | 1.9 (1.5) |
| **Paroxysmal AF [%]** | 24 [57] |
| *Echocardiographic Parameters* | |
| **LVEF [%]** | 57.5 (10.0) |
| **LAVi [ml/m$^2$]** | 42.9 (8.5) |
| **3D LA min vol [ml/m$^2$]** | 43.4 (20.2) |
| **3D LA max vol [ml/m$^2$]** | 68.2 (20.4) |

Supplemental Table 3: HD-grid-mapped AF Patient Characteristics

#### S2 – COMPUTER SIMULATED AF

The Courtemanche model of AF used here was adapted from one previously described.[1] The Courtemanche model of the human atrial cardiomyocyte was used,[2] that was adapted and implemented in as a monodomain model in CARP, running in a parallel cloud-based cluster. Tissue conductance was set to provide physiological anisotropy with conduction velocity in a longitudinal direction of 47.9 cm/s. The tissue slab was 7x6cm in size, with longitudinal fiber anisotropy with grid ration of 6. Grid discretization was performed at 100μm resolution. The membrane capacitance was set to 1μF/cm.[2] Differential equations were solved with a 25μs timestep, with simulations of up to 5 seconds. Atrial fibrillation epochs were initiated by a standard S1-S2 cross-shock protocol. 2D simulations were performed with a ACh distribution was generated by randomly assigning a value between 0 and 0.001 μM to each node (cardiomyocyte). The ACh-activated $K^+$ current was implemented using the previously-published model by Kneller et al.[3] AF epoch transmembrane voltage movies were visualized in Meshalyzer (OpenCARP), and exported as transmembrane voltage movies.

#### S3 – DATA PRE-PROCESSING

To allow for phase and wavefront detection, electrode coordinates were projected on a 2D plane.[4, 5] The three-dimensional co-ordinates of mesh vertices were mapped onto a 2D polar plot using a cone-shaped surface projection and Delaunay triangulation. Using this 2D projection, electrode potentials were linearly interpolated from the electrodes onto a fine regular grid (100x100 grid points).

To remove electrodes possessing poor signal-to-noise ratios, these electrograms were removed from the analyses prior to 2D projection of the mesh by selecting only signals with a dominant frequency within the 1.5 to 45 Hz band for analyses. EGMs near the stimulus electrode at times also demonstrated an exponential artifact, as well as respiration artifacts. To remove this, a mode was fitted with a constant offset of the signal mean, an exponential term, as well as a sinusoidal term. This fitted signal was subtracted from the raw signal to improve signal to noise ratio.

### S4 – RENEWAL THEORY

#### S4.1 – Renewal rate constants

The average rate of spiral formation (given by $\lambda_f$) can be measured by calculating the exponential decay constant from the inter-formation time distribution:

$$f(t) = \left\{\lambda_f e^{-\lambda_f t} \quad t \geq 0\right\} \quad (3)$$

where $t$ is time, and $\lambda_f$ the formation rate.[22] Similarly, the average rate of destruction (given by $\lambda_d$) is given by:

$$f(t) = \left\{\lambda_d e^{-\lambda_d t} \quad t \geq 0\right\} \quad (4)$$

where $t$ is time, and $\lambda_d$ the destruction rate.

The interaction of $\lambda_f$ and $\lambda_d$ is given by the '*mixing rate*', which represents the time taken to reach the stationary state distribution. The mixing rate is specifically given by[24]:

$$Mixing\ rate = \log(1 - z) \quad (5)$$

where $z$ is the second largest eigenvalue modulus (or SLEM) of the Markov transition matrix described by $\lambda_f$ and $\lambda_d$ (further detail provided in **Supplement S4.2**). In this study, $\lambda_f$ and $\lambda_d$ were calculated as previously described from pre-processed and Hilbert transformed unipolar electrograms, with the PS detection algorithm further detailed **Supplement S5**.[20,22]

#### S4.2 – The M/M/∞ Markov birth-death transition matrix

The following section is a brief illustration of the M/M/∞ birth-death transition matrix adapted from Crawford, 2018.[6] As mentioned previously, birth-death processes count the population in a system at a time $t$. For an M/M/∞ birth-death process, we consider the behavior of the system by considering a short time $dt$ when there are $k$ particles (in our case PS, or WF in the system). As $dt$ approaches zero, the probability of an event in the interval $(t, t + dt)$, given $X(t) = k$, and $\mu_k = k\mu_1$:

$$P_{ab}(t + dt) = \lambda_{b-1}P_{a,b-1}(t)dt + \mu_{b+1}P_{a,b+1}(t)dt + (1 - \lambda_b - \mu_b)P_{ab}(t)dt + o(dt)$$

$$P = \begin{bmatrix} P_{i,j} & P_{i,j+1} \\ P_{i+1,j} & P_{i+1,j+1} \end{bmatrix}, \quad P_{i,j} = \begin{cases} p_i, & if\ j = i + 1 \\ q_i, & if\ j = i - 1 \\ 1 - p_i - q, & if\ j = 1 \\ 0, & else \end{cases}$$

$$P = \begin{bmatrix} P_{0,0} & P_{0,1} & P_{0,2} & P_{0,3} & P_{0,4} & P_{0,5} & P_{0,6} \\ P_{1,0} & P_{1,1} & P_{1,2} & P_{1,3} & P_{1,4} & P_{1,5} & P_{1,6} \\ P_{2,0} & P_{2,1} & P_{2,2} & P_{2,3} & P_{2,4} & P_{2,5} & P_{2,6} \\ P_{3,0} & P_{3,1} & P_{3,2} & P_{3,3} & P_{3,4} & P_{3,5} & P_{3,6} \\ P_{4,0} & P_{4,1} & P_{4,2} & P_{4,3} & P_{4,4} & P_{4,5} & P_{4,6} \\ P_{5,0} & P_{5,1} & P_{5,2} & P_{5,3} & P_{5,4} & P_{5,5} & P_{5,6} \\ P_{6,0} & P_{6,1} & P_{6,2} & P_{6,3} & P_{6,4} & P_{6,5} & P_{6,6} \end{bmatrix}$$

$$P = \begin{bmatrix} 1 - p_i - q_i & p_i & 0 & 0 & 0 & 0 & 0 \\ q_i & 1 - p_i - q_i & p_i & 0 & 0 & 0 & 0 \\ 0 & q_i & 1 - p_i - q_i & p_i & 0 & 0 & 0 \\ 0 & 0 & q_i & 1 - p_i - q_i & p_i & 0 & 0 \\ 0 & 0 & 0 & q_i & 1 - p_i - q_i & p_i & 0 \\ 0 & 0 & 0 & 0 & q_i & 1 - p_i - q_i & p_i \\ 0 & 0 & 0 & 0 & 0 & q_i & 1 - p_i - q_{i.} \dots \end{bmatrix}$$

#### S4.3 Eigenvalue spectrum, spectral gap and mixing rates of a Markov birth-death process

The transition matrix of a Markov birth-death process is often of interest as it can be used to study the long-term population distribution of a system. Mathematically speaking, this stationary state distribution can be identified by finding the eigenvector which has at least one eigenvalue ($\lambda$) = 1. In simple terms, an eigenvector is a vector that does not change in direction when a linear transformation is applied to it, whilst eigenvalues are the scalar quantities that define how the eigenvector is scaled due to the given linear transformation.

Studying the eigenvalues for a given transition matrix, i.e. the *'eigenvalue spectrum'*, can give insights into the dynamics of that system. For example, the difference between the first and second largest eigenvalues is referred to as the '*spectral gap*' and describes how quickly the steady state distribution is reached. The spectral gap is given by[7]:

$$1 - z$$

where 1 is the largest eigenvalue and $z$ the second largest eigenvalue modulus (often referred to as SLEM).

The time taken to reach the stationary state distribution can also be expressed in terms of the '*mixing rate*' of the birth-death process, given by:

$$\log(1 - z)$$

If SLEM is very close to 1, then the mixing rate will approximate the spectral gap.

In this study, we specifically hypothesized that AF termination would occur due to a deviation from the steady state of AF dynamics, due to the steady state distribution being reached more slowly. This consequently would provide a greater opportunity for the process to diverge from the steady state distribution and break the cycle of PS and wavelet regeneration.

#### S5 – PHASE SINGULARITY AND WAVEFRONT DETECTION AND TRACKING

In this study, we used a phase singularity (PS) detection algorithm based on a convolution kernel method.[6, 7] The convolution kernel method approximates the gradient of phase from the discretized phase map using a finite difference operation in the $x$ and $y$ directions, given by:

$$k_x[m,n] = \nabla_{\phi_x}[m,n] = \phi[m+1,\ n] - \phi[m,\ n]$$

$$k_y[m,n] = \nabla_{\phi_y}[m,n] = \phi[m,\ n+1] - \phi[m,\ n]$$

where is $\phi$ phase, and $[m,n]$ the pixel coordinates. The line integral at the pixel $[m,n]$ can there be approximated using the convolution kernel equation:

$$\left(\nabla X\vec{k}\right)\cdot\hat{z} \propto \nabla_x \otimes k_y + \nabla_y \otimes k_x$$

where $\otimes$ is the convolution operator, and $\nabla_x$ and $\nabla_y$ and convolutional kernels.

In order to track PS, a tracking algorithm was implemented as previously described. New PS were defined as the detection of a PS not falling within the surrounding radius of $r$ of an existing PS for a duration of $\tau$. As default, $\tau$ was set to 10ms (20 frames) and $r$ set to account for the spatial resolution of the mapped field.

To determine PS lifetime and inter-formation times, a look-up table indexing onset time, offset time and electrode location for each PS was created. If the PS does not fall within a radius $r$ of another, the P is given a new ID by incrementing the list of current PS IDs by 1. The look-up table will then record the current time stamp as the first time the PS is detected, and records all consequent time stamps it remains present. The look-up table also records the x and y locations of the PS on the phase map. For PS, this is given by a paired (x,y) coordinate value. Using this look-up table information, the total time the PS is present (lifetime) can be measured, as well as the time taken between the creation of the current PS until the creation of the next consecutive PS (inter-formation time).

#### S6– STATISTICAL ANALYSIS

Differences in fractal dimension, correlation length, $\lambda_f$, $\lambda_d$ and mixing rates between cases of sustained and spontaneously terminating AF, VF, and computer simulated fibrillation was assessed using generalized linear mixed models with gamma distribution, with $P<0.05$ indicating significant differences. Similarly, differences in fractal dimension and correlation length between i) paroxysmal and persistent AF episodes were also assessed using a gamma generalized linear mixed model (significance at $P<0.05$).

Fractal dimension and correlation length were also measured through different stages of perfused VF, ischemia and reflow to assess whether any changes could be observed during VF progression. Specifically, a generalized linear mixed effects models was used to study changes in either fractal dimension or correlation length (set as the target variable) with stage (perfusion, ischemia and reflow) set as the fixed effect.

A similar approach was also used to study the changes in fractal dimension and correlation length over time post flecainide administration, with time segment (broken into 5-minute intervals) set as the fixed effect. A paired Wilcoxon's signed rank test was additionally used to observed whether differences in pre vs. post flecainide administration epochs from the same patient were statistically significant (significance at $P<0.05$).

To study the relationship between $\lambda_f$ and $\lambda_d$ with fractal dimension and correlation length, a k-means clustering approach was implemented using 10 replicates and a city-block algorithm as the distance metric.

#### S7– EXTENDED DATA FIGURE 1

Human ventricular fibrillation episodes from n=12 patients undergoing cardiac surgery were studied. Three distinct stages of VF were present during the observed recordings: i) the perfusion stage (initial 30 seconds); ii) ischemia (following 150 seconds); and iii) reflow (last 30 seconds). **Extended Data Fig.1** depicts the propagation of electrical activity as VF evolves throughout the three stages, which can be seen to depict more turbulent behaviour as VF progresses. This evolution resulted in changes to the correlation length, which decreased as the VF dynamics became more turbulent. The fractal dimension conversely became larger as VF evolved.

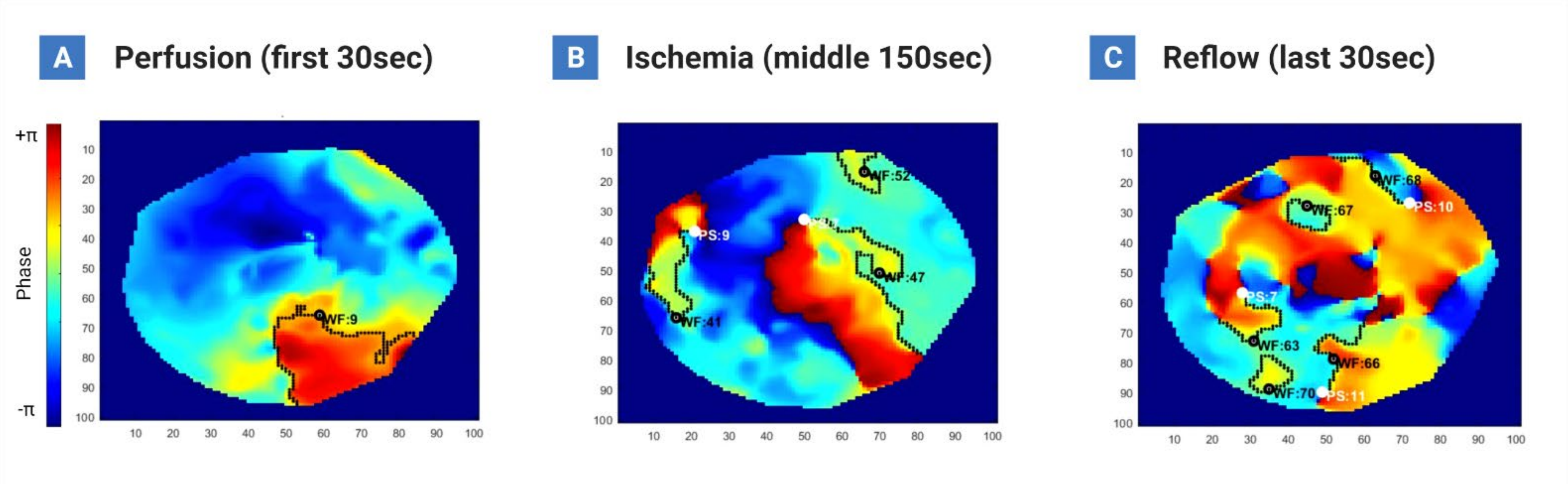


**Extended data Fig. 1**
